# Dual-color single molecule localization microscopy on transparent polymer waveguide chips

**DOI:** 10.1101/2022.11.29.518375

**Authors:** Anders Kokkvoll Engdahl, Surjendu Bikash Dutta, Stefan Belle, Jasmin Schürstedt, Karolina Szafranska, Peter McCourt, Ralf Hellmann, Thomas Huser, Mark Schüttpelz

## Abstract

Photonic waveguide chips offer near-field excitation of biological samples, which enables cost-effective, large field-of-view super-resolution microscopy without the need for high numerical aperture (NA) objective lenses. Single molecule localization based super-resolution microscopy that requires high illumination intensities is currently limited to solid state photonic waveguide chips composed of hard-coated, high NA planar waveguides deposited on opaque substrates. These platforms do not permit epi-detection of fluorescence through the substrate, which limits the use of photonic waveguide chips to the upright configuration. Additionally, the detection efficiency is reduced because the majority of the fluorescence emission is directed towards the high refractive index substrate. A low cost waveguide chip based on a polymer core material deposited on common #1.5 coverslips that is easy to produce was recently demonstrated. Here, a platform that is capable of performing single-molecule localization microscopy (SMLM) of biological samples using polymer-based photonic waveguide chips is presented, enabling super-solution microscopy in the inverted microscope configuration. Super-resolved imaging of two different structures of the cytoskeleton in primary liver sinusoidal endothelial cells (LSECs) by two popular SMLM methods, *d*STORM and DNA-PAINT, down to 23 nm is demonstrated.

## 1 Introduction

In photonic waveguide chip-based microscopy the sample is illuminated through planar waveguides with a high refractive index [1]. The evanescent field created by total-internal reflection (TIR) at the interface between the waveguide and the sample can generally be used for both fluorescence [2, 3] and light scattering [4, 5] microscopy, but decays exponentially with a typical penetration depth in the order of ∼0.1 µm into the sample. High light intensities at the waveguide-to-sample interface, which rapidly decay with increasing distance from the waveguide surface, provide waveguide chips with axially super-resolved sample excitation. Since fluorophores that are further away from the surface of the waveguide are not excited by the evanescent field, out-of-focus fluorescence is reduced to a minimum. This makes waveguide-based total-internal reflection fluorescence (TIRF) excitation ideal for single-molecule localization microscopy (SMLM) techniques, such as (*d*)STORM [6–10] and (DNA-)PAINT [11–13], where a high signal-to-noise ratio is required for the precise localization of single emitters with low uncertainty. An important limitation for the wide-spread use of this technique, however, has been the complex manufacturing process of such waveguide chips. Photonic waveguide chips for SMLM have, until now, exclusively been manufactured on opaque substrates made of thick silicon wafers. This requires advanced techniques for the deposition of core and cladding layers, followed by electron-beam and ion-beam lithographic techniques to shape the individual waveguides within the deposited waveguide layer. These methods require access to expensive clean-room fabrication facilities, which lead to rather high production costs of ∼100 EUR per chip [10]. Additionally, due to the opacity of the material they can only be used in the upright detection configuration, which leads to optical aberrations and limits biological applications, such as imaging of adherent cells. Efforts have been made to produce thin transparent chips based on quartz glass [14]. In addition to a further increase in production costs, the brittle nature of this material makes handling of such chips extremely delicate, again limiting their widespread use.

Recently, we introduced an affordable and highly efficient transparent photonic waveguide chip based on the negative photoresist polymer EpoCore which is spin-coated on top of ordinary #1.5 coverslips [15, 16]. Transparent polymer waveguide chips used in combination with an inverted microscopes enabled waveguide-TIRF imaging of biological samples with an unprecedented large field-of-view (FOV). Even super-resolution imaging was demonstrated with polymer waveguide chips by post-processing of the waveguide-TIRF based images exploiting super-resolution radial fluctuations (SRRF) [17]. Although SRRF is a versatile method, it is prone to generate imaging artefacts arising from high-density labeling and images with low fidelity [18]. Here, we present the application of enhanced polymer-based transparent waveguide chips on a modified imaging platform to enable large FOV imaging with superior and robust super-resolution microscopy methods. We show that these chips are well suited for SMLM techniques that typically require rather high illumination intensities, such as *d*STORM and DNA-PAINT. Because the detection of fluorescence emission is decoupled from the fluorescence excitation on waveguide chips, a non-TIRF objective lens can be used for detection. Hence, SMLM on these transparent waveguide chips can even be performed with an inexpensive microscope configuration that makes use of a low cost industry-grade CMOS camera for fluorescence detection. We demonstrate this by multi-color super-resolution imaging of different fluorescently labeled cytoskeletal structures in primary liver sinusoidal endothelial cells (LSECs).

## 2 Experimental section

### 2.1 Setup

The main part of the microscope consists of a 3D-printed chip holder mounted on an XYZ-axis positioning stage (M-562-XYZ, Newport) and a beam XYZ-axis piezo positioning stage (Nanomax 300, Thorlabs) for coupling the beam into the waveguide end facet (**Figure 1a**). Since coupling into the multi-moded waveguide produces specific interference patterns causing intensity fluctuations and stripe patterns we gradually vary the position of the laser beam at the waveguide end facet over time using an open-loop piezo controller (MDT693b, Thorlabs). Applying a sine function with an amplitude of ∼50 V and a frequency of ∼0.3 Hz) to the x-axis stage position (along the chip end facets) results in a lateral amplitude of ∼13 µm such that stripe patterns in SMLM raw data images are averaged out. In order to use the waveguide chips with high resolution objective lenses we designed a 3D-printed chip holder that holds the chip firmly without limiting the vertical position of the objective lens relative to the waveguide chip (**Figure 1b**). The waveguide chip is inserted into the holder and held in place by narrow clamps on the sides. The chip holder is attached to a chip-to-stage adapter that is made of aluminium for stability reasons (**Figure 1c**). As laser source we use an Ar-Kr^+^ ion laser (Innova 70C Spectrum, Coherent) emitting light with a wavelength of 568 nm and a power of ∼50 mW, and 647 nm with ∼150 mW. SMLM raw data images are acquired with an uncooled industry-grade CMOS camera (UI-3060CP-M-GL, IDS Imaging Development Systems) with an 1936 × 1216 pixel sensor and a pixel size of 5.86 µm. For fluorescence collection we use a 60x 1.35 NA oil immersion objective lens (UPlanSApo, Olympus), a plano-corrected objective lens with high NA, which is too low for objective-type TIRF microscopy.

**Figure 1:**
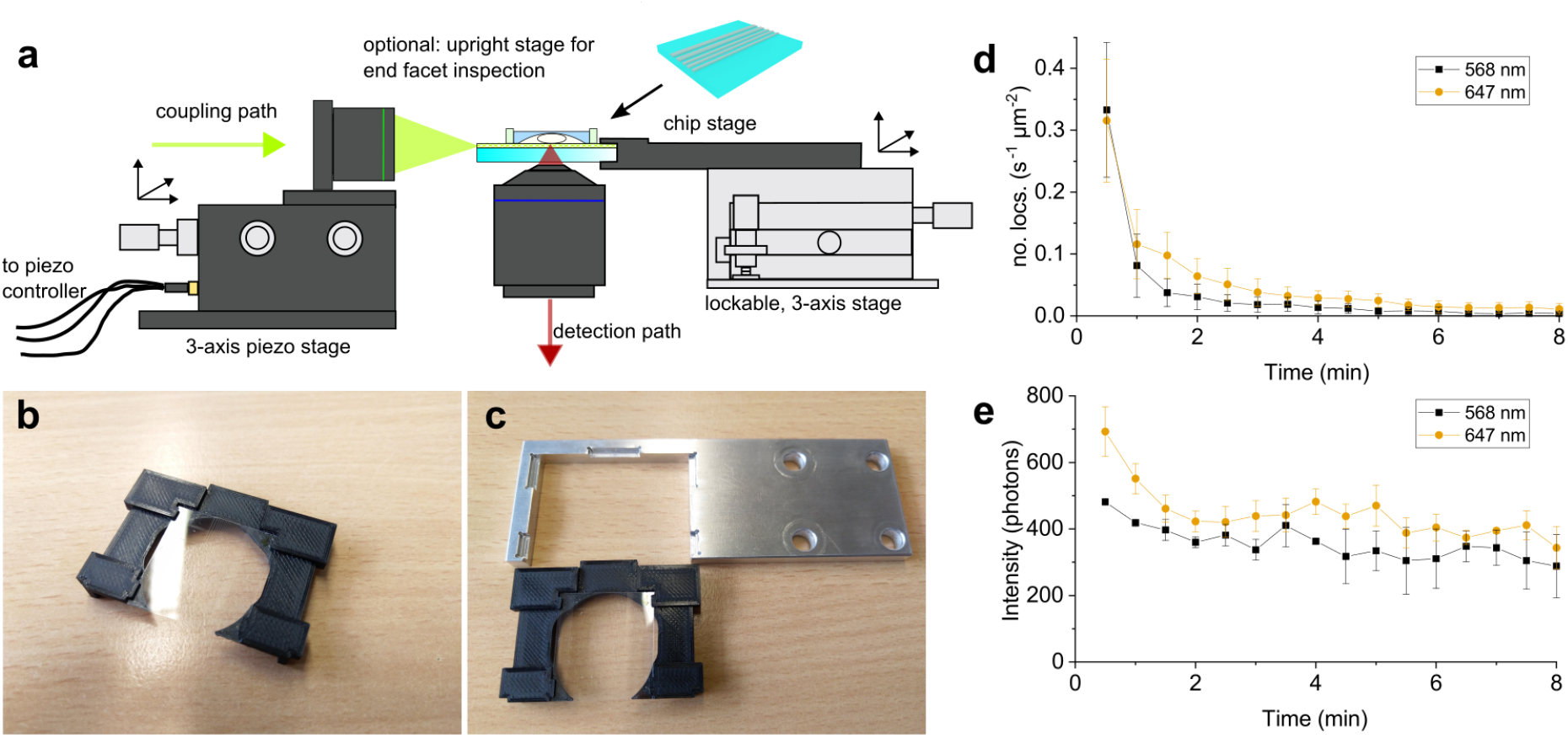
**a)** Schematics of the waveguide chip microscope. The main part of the microscope consists of two three-axis stages. An XYZ-axis piezo positioning stage is used to couple excitation light into the chip, while another XYZ-stage is used to hold and position the chip with respect to the detection optics. **b)** The 3D-printed chip holder keeps the chip firmly in place and protects it from mechanical stress. **c)** The chip holder can easily be clicked in or removed from the aluminium chip-to-stage adapter. **d)** The number of localizations of the autofluorescent background of the waveguide decreases during the first few minutes due to photobleaching. The number of emitters in the polymer layer drastically drops if high illumination powers are coupled into the layer. **e)** The intensity of autofluorescent background emitters in the polymer layer decreases accordingly and converges to ∼400 photons for both excitation wavelengths.

### 2.2 Sample preparation

#### 2.2.1 Chip preparation

We produce EpoCore waveguide chips with 1.2 µm core height as reported earlier [15]. The chips are first visually inspected by coupling the beam into a set of waveguides, then cleaned by submersion in a ultrasonicator bath containing 0.05% sodium dodecyl sulfate (SDS) for 5 minutes. Following a rinsing step in water and ethanol, the chips are then dried with N_2_ and a PDMS sample chamber is placed on top.

#### 2.2.2 Cell seeding

Before seeding cells on the waveguide chip the sample chamber surface is coated with fibronectin (0.2 mg ml^*−*1^ in Phosphate Buffered Saline (PBS) containing 2 mM ethylenediaminetetraacetic acid (EDTA)) for 1 hour at room temperature and washed with PBS afterwards. Cryo-preserved LSECs from rats are stored at −80 °C. For thawing and seeding, the vial with the LSECs is placed in the incubator at 37 °C until nearly all ice is thawed. The cells are gently pipetted drop-wise to 25 ml of pre-warmed Dulbecco’s Modified Eagle Medium (DMEM) and centrifuged at 50 g for 3 minutes to remove other cell types, e.g. hepatocytes. The supernatant containing the LSECs is used for a second centrifugation step at 300 g for 8 minutes. The cell pellet is resuspended in 4 mL to 7 mL DMEM and 200 µl (∼100.000 cells per cm^2^) of the cell solution is added to the coated waveguide. After 1 hour of incubation at 37 °C in the incubator, the cells are washed with pre-warmed DMEM and incubated for another 2 hours before fixation.

#### 2.2.3 Sample staining

For indirect antibody-staining of tubulin filaments, the liver sinusoidal endothelial cells are fixed according to [19]. An extraction step is followed by fixation with 0.5% glutaraldehyde in PEM buffer (PEM: 80 mM piperazine-N,N-′bis(2-ethanesulfonic acid) (PIPES), 5 mM egtazic acid (EGTA), 2 mM MgCl_2_ at pH 6.8). After washing with PBS, glutaraldehyde-induced autofluorescence is quenched by addition of 0.1% NaBH_4_ in PBS for 7 minutes followed by washing with PBS three times. Cells are blocked and permeabilized using a blocking buffer containing 0.3% gelatin and 0.05% Triton X-100 in PBS for 1 hour. Staining is performed overnight at 4 °C for *α*- and *β*-tubulin using a mixture of three primary antibodies (T5168, T6199, T5923, Sigma) at a combined dilution of 1:150 in blocking buffer. Cells are washed with PBS three times and, for *d*STORM experiments, the second staining solution, AF647-conjugated goat anti-mouse IgG secondary antibody (A-21237, ThermoFisher) diluted 1:200 in blocking buffer, is incubated at room temperature for 90 minutes. Alternatively, the same staining step is used with a donkey anti-mouse IgG antibody conjugated with docking strand 1 (Massive-AB 1-Plex, Massive Photonics) when staining for DNA-PAINT. Actin staining using phalloidin-AF647 is performed either after or during the secondary antibody-staining step at a 1:40 dilution in blocking buffer. The cells are washed 5 times with PBS before postfixation (3 minutes with 1% paraformaldehyde in PBS). For DNA-PAINT and dual color experiments, we add 0.1 µm fluorescent TetraSpeck beads (T7279, Invitrogen) as drift markers to the sample (at 1:500 dilution in PBS, left to settle until a sufficient number of beads is detectable).

### 2.3 SMLM experiments

#### 2.3.1 Reconstructions

A super-resolution image is reconstructed from the series of acquired raw data images with the SMAP [20] package using a basic fitting model and correcting for pixel gain and base level variations of the CMOS camera sensor. The camera sensor was characterized beforehand by a photon-free method and the plugin ACCéNT [21] for *µ*Manager [22]. The localization tables are then imported into the ImageJ-plugin ThunderSTORM [23, 24] and visualized as average shifted histograms with 5× or 10× image dimensions.

#### 2.3.2 *d*STORM buffers

Depending on the width of the waveguides and the fluorophore we use different buffers and switching agents in order to achieve uniform blinking of the fluorophores when imaging in waveguide-TIRF. For narrower waveguides we use the common GODCAT buffer containing the enzymatic oxygen scavengers, glucose oxidase and catalase, with beta-mercaptoethanol (BME) as switching agent. For wider waveguides we use the recently reported sulfite buffer [25], containing 1-90% glycerol and 10 mM to 50 mM Na_2_SO_3_ in PBS. For lower glycerol concentrations it is preferable to add either p-phenylenediamine (PPD) (100 mg L^*−*1^) or cyclooctatetraene (COT) (2 mM) as an extra preservative of on-state fluorophores. Sulfite buffers without GODCAT are usually highly effective for switching AF647 but come at the cost of lower fluorophore brightness when compared to 143 mM BME in GODCAT.

## 3 Results

### 3.1 Polymer autofluorescence

We first measured the autofluorescence in pure 1.2 µm tall EpoCore waveguides. Autofluorescent spots in the polymer layers arising from impurities can switch on and off similar to fluorophores in SMLM measurements, which leads to an erroneous background contribution in super-resolved images. After cleaning the substrate, we applied PDMS chambers to the substrate that were filled with double-distilled water. Laser light with a wavelength of 568 nm or 647 nm and a power of ∼25 mW was coupled into the 120 µm wide waveguides and fluorescence was measured over 15 minutes using 50 ms camera exposure per frame. The mean intensity values from three separate measurements was plotted and shows that the number of autofluorescent emitters rapidly decreased down from ∼0.3 s^*−*1^ µm^*−*2^ to a level of ∼0.01 s^*−*1^ µm^*−*2^ during the first few minutes of irradiation (**Figure 1d,e**). The localization intensity also decreased from a starting value of 700 photons and 500 photons for 568 nm and 647 nm excitation, respectively, to ∼400 photons for both wavelengths. As we saw for bulk autofluorescence in [15], we expect the autofluorescent impurities to slowly recover over the course of multiple days. This shows that autofluorescence in the polymer is readily suppressed after the first few minutes of any SMLM experiment and will not lead to artefacts in the reconstructed images.

### 3.2 SMLM of the LSEC cytoskeleton

We next proceeded to attempt SMLM experiments on fixed biological samples. **Figure 2** shows *d*STORM experiments on Alexa Fluor 647 (AF647) stained tubulin filaments in rat liver sinusoidal endothelial cells adhering to a 50 µm wide polymer waveguide. Due to the absence of an evanescent field near the edges of the waveguide, these regions appear as dark areas at the top and bottom of the images. Here, we used an enzymatic *d*STORM buffer containing glucose oxidase and glucose catalase (GODCAT) to remove oxygen, combined with 143 mM beta-mercaptoethanol (BME) to cause intermittent fluorescence emission of AF647. 40,000 frames were acquired at 50 ms exposure time per frame using 647 nm laser light with a power of ∼35 mW measured at the waveguide end facet. The laser spot was rapidly swept across the waveguide facet to average out multi-mode intensity patterns within the waveguide. 200 images at ∼1 mW were acquired to obtain an average widefield image, in which the stripe pattern is still visible, while this pattern is not visible in the reconstructed *d*STORM image (**Figure 2a,b**). This becomes even more apparent when a tubulin filament is visible in the *d*STORM reconstruction (**Figure 2e**) that is not clearly visible in the corresponding widefield image (**Figure 2c**). To estimate the spatial resolution of our polymer waveguide-chip based SMLM experiments, the Fourier Ring Correlation (FRC) analysis was applied to these data [26, 27]. The corresponding FRC plots are shown in **Figure 2g** and reveal a resolution of 23 nm and 25 nm for the filaments shown in the insets in **Figure 2e** and **f**, respectively. Further analysis of 6.8 million emitter localizations in the *d*STORM image yields histograms of the intensity distribution, background standard deviation and localization uncertainty per localized emitter **Figure 2h**. A median average of 725 photons for the localization intensity and 10.9 photons for the background standard deviation demonstrate that these data exhibit an excellent signal-to-noise ratio that leads to a localization uncertainty as low as 12.0 nm.

**Figure 2:**
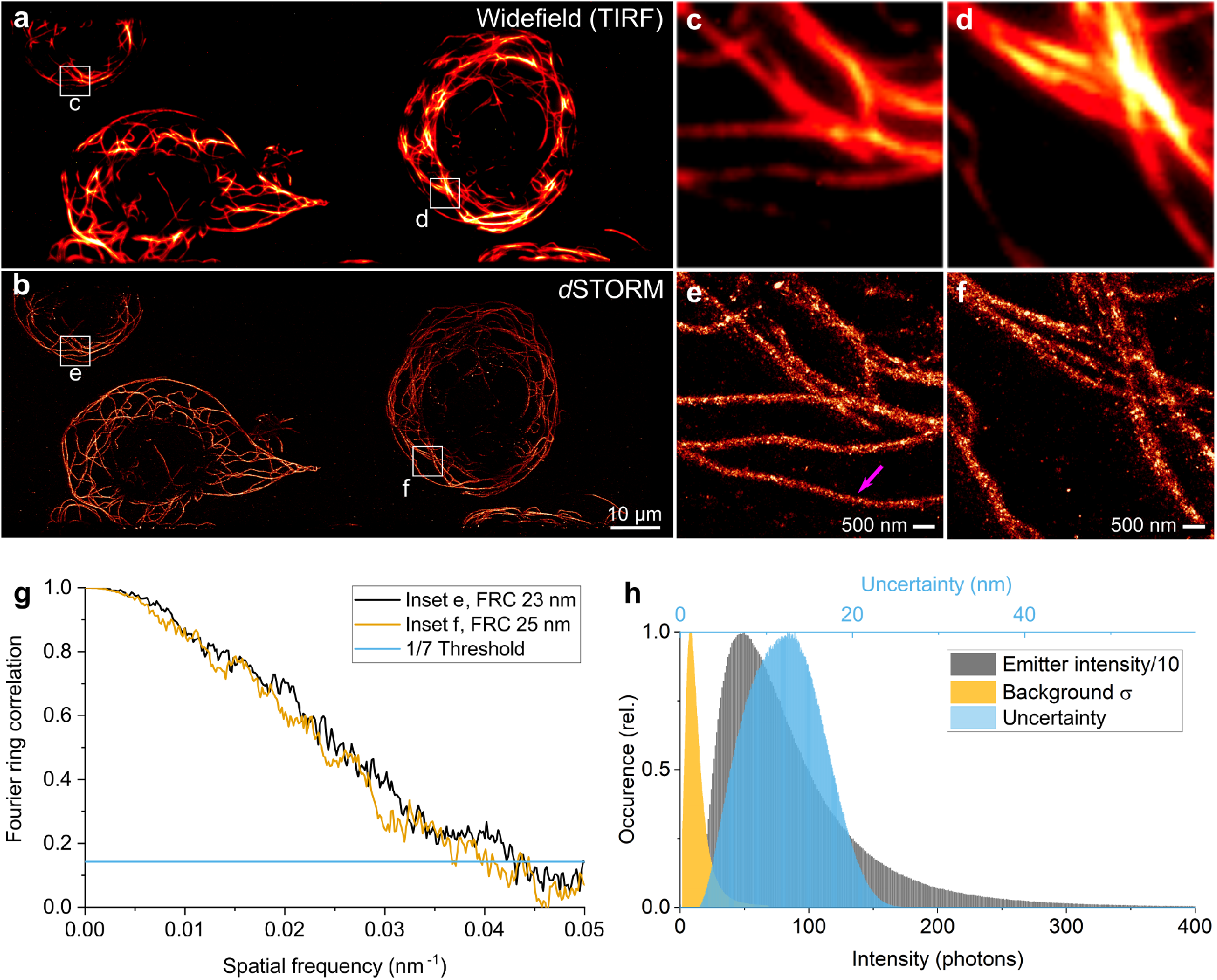
**a)** Widefield image of Alexa Fluor 647-stained tubulin filaments in fixed rat liver sinusoidal endothelial cells plated on a 50 µm wide polymer-based photonic waveguide. The average intensity of 200 frames was recorded at low laser power (∼1 mW) over 10 s in order to average out multi-mode intensity patterns in the waveguide. **b)** *d*STORM image of the super-resolved tubulin cytoskeleton reconstructed from 40,000 frames recorded at high laser power (∼35 mW). **c**,**d)** The insets provide magnified widefield images of the region of interest marked with white boxes in **a. e**,**f)** The same region of interest is chosen for reconstruction of super-resolved *d*STORM images. A tubulin filament is visible in the *d*STORM reconstruction (**e**, marked with an arrow) that is not clearly visible in the corresponding widefield image **c. g)** The spatial resolution of the insets is calculated by Fourier Ring Correlation (FRC). The threshold (1/7) is used to determine the spatial resolution of 23 nm in **e** and 25 nm in **f**, respectively. **h)** Histograms of the intensity, background standard deviation and localization uncertainty per localization of the *d*STORM image in **b**.

We also successfully performed DNA-PAINT measurements on polymer-based photonic waveguides. In DNA-PAINT, the structure of interest is first identified using an antibody linked to a target DNA strand. A fluorescently labeled imager DNA strand is then added, which transiently binds to the target strand and enables localization of the attached fluorophore during this short time of immobilization. Here, we used an imaging buffer containing 100 pM ATTO565-conjugated imager DNA strands. Excitation laser light with a wavelength of 568 nm and a power of 50 mW at the end facet of the waveguide was used. Here, we acquired 30,000 frames using 200 ms camera exposure time per frame. **Figure 3** shows a DNA-PAINT reconstruction of immunostained microtubules on a 120 µm wide waveguide. The reconstruction demonstrates the low affinity of the fluorescently-labeled imager strand to the polymer layer resulting in a low background signal. This underlines the broad applicability of polymer waveguide chips for different SMLM techniques - additionally enabling a large field-of-view of ∼120 µm ×200 µm for super-resolved images. Generally, both SMLM techniques, *d*STORM and DNA-PAINT, can be used in the same experiment, i.e. for multi-color staining of different biological targets. We used this approach for a two-color staining of the actin and the tubulin cytoskeleton of rat LSECs. Actin filaments were labeled with phalloidin-AF647 and imaged with *d*STORM using an imaging buffer containing glycerol and sulfite [25]. Tubulin was imaged using 100 pM of an ATTO565-conjugated imager strand in an imaging buffer containing 5% glycerol. *d*STORM and DNA-PAINT raw data images were acquired sequentially using 647 nm and 568 nm for excitation, respectively. Camera exposures were 50 ms and 100 ms. **Figure 4** shows the reconstruction of 20,000 frames acquired for each color using ∼20 mW to 30 mW laser power on a 60 µm wide waveguide. The field-of-view acquired with the camera is 60 µm × 150 µm and shows an overview image of three different cells on the waveguide. Magnified insets shown in **Figure 4b-d** reveal the super-resolved actin and tubulin cytoskeleton of these cells. Both techniques, although applied to the biological sample in the same experiment, did not negatively impact each other through e.g. unspecific bindings. Also the density of localizations is ∼50 times higher on cellular targets than on the pure waveguide demonstrating the high specificity of SMLM methods on the EpoCore waveguides.

**Figure 3:**
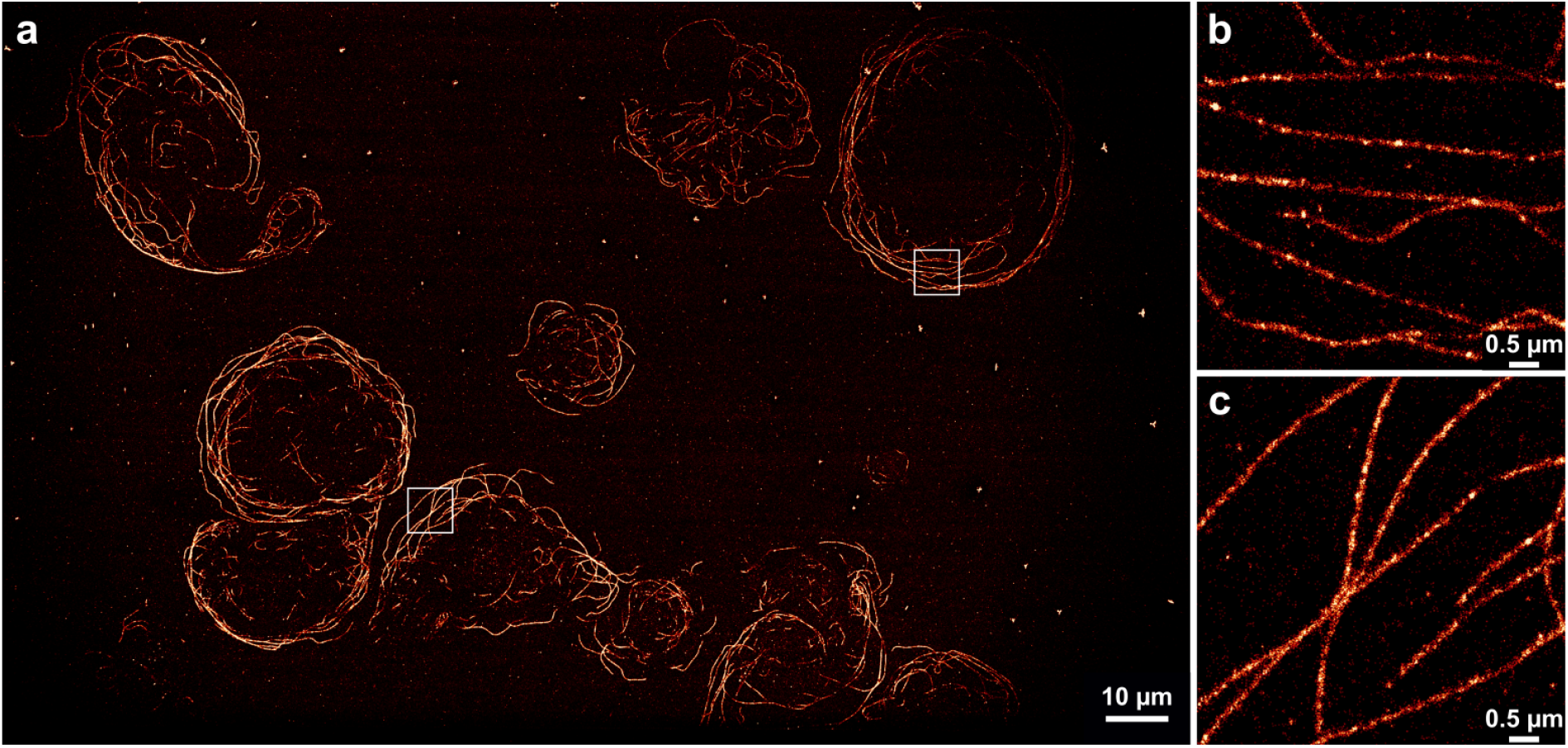
**a)** Wide field-of-view DNA-PAINT reconstruction of tubulin filaments labeled with dockingstrand-conjugated antibodies in fixed rat liver sinusoidal endothelial cells on a 120 µm wide EpoCore waveguide. 20,000 frames recorded at ∼50 mW laser power using a 568 nm laser were used for the reconstruction of the image. **b**,**c)** The insets provide magnified views of the regions of interest marked with white boxes in **a**.

**Figure 4:**
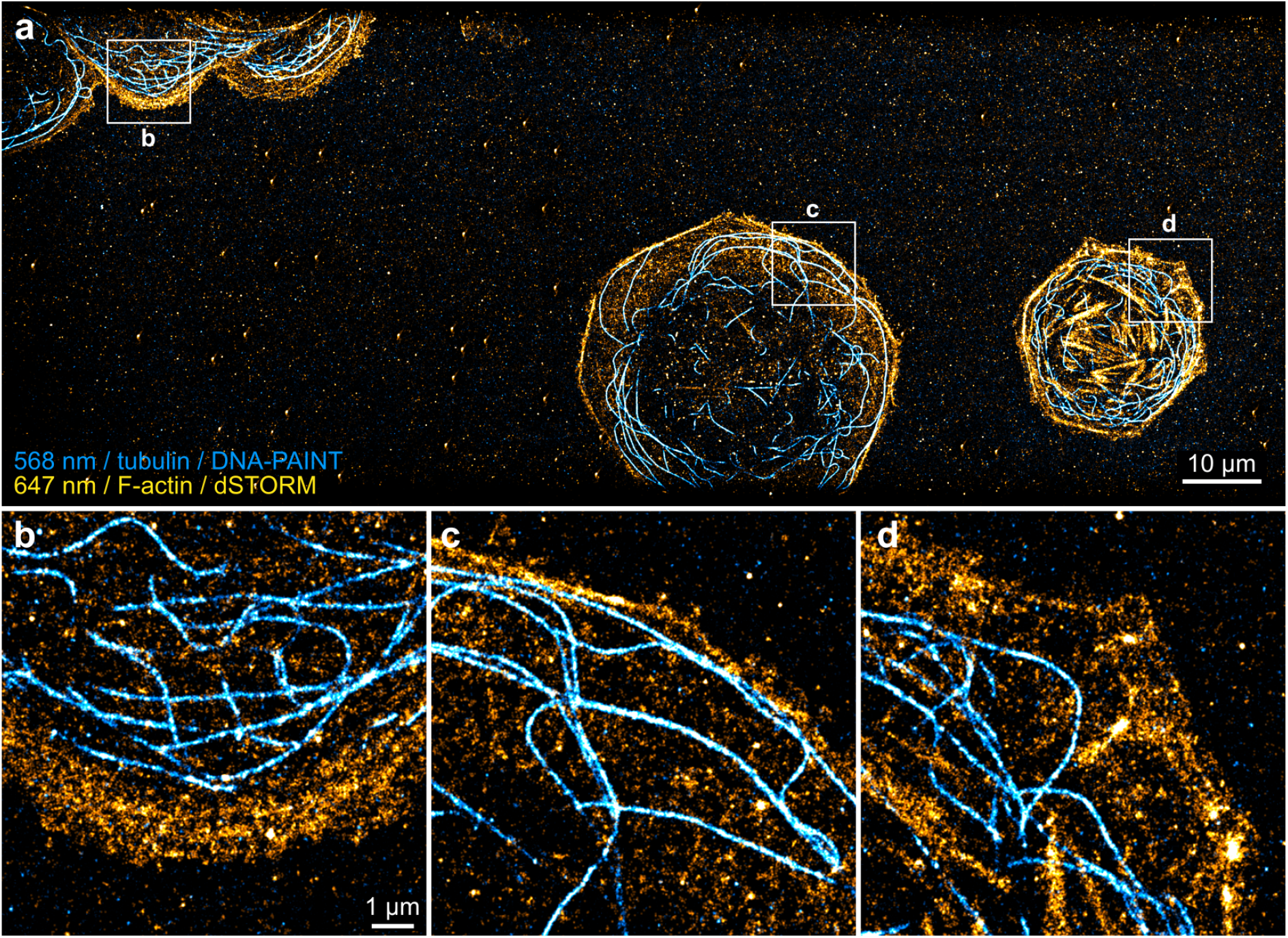
**a)** Two-color SMLM reconstruction of actin (yellow) and tubulin (cyan) in rat liver sinusoidal endothelial cells plated on a 60 µm wide waveguide. **b-d)** Magnified views of the region of interest marked with white boxes in **a**. Actin fibers were stained with Alexa Fluor 647-phalloidin and imaged using *d*STORM with a wavelenght of 647 nm and tubulin filaments were stained with anti-tubulin antibodies and imaged using DNA-PAINT with ATTO565 using 568 nm for excitation.

## 4 Conclusion

In this work we have shown that polymer-based photonic waveguide chips offer illumination intensities and detection sensitivity for two different single-molecule localization microscopy modalities, i.e. *d*STORM and DNA-PAINT, in biological samples. To the best of our knowledge, this is the first time that these SMLM techniques have been demonstrated on transparent polymer waveguide chips in the inverted microscope configuration. Autofluorescent single emitters of the polymer layer are rapidly photobleached during the first minutes of imaging. They can be further suppressed by prior photobleaching procedures so that they do not contribute to the background during image acquisition. Polymer waveguides deposited on standard #1.5 coverslips are much easier to produce and at a fraction of the cost (∼1%) when compared to the rather expensive manufacturing process of hard-coated, silicon wafer based waveguide chips. The thin coverslips allow us to take advantage of the highest resolution objective lenses and avoid aberrations that arise from imaging through the sample chamber in the upright mode. By using an industry-grade CMOS camera for detection, we could show that super-resolution imaging with a resolution of 23 nm is possible at a very low cost, while still achieving large field-of-views.

## Supporting information

3D CAD archive for chip holders

## Supporting Information

CAD files for the chip holder and the chip-to-stage adapter are available for download as supporting information to this article.

## Acknowledgements

M.S. acknowledges funding from the European Regional Development Fund (ERDF), grant no. EFRE-0400137. P.McC., K.S., and T.H. were supported by the European Union’s Horizon 2020 research and innovation program under the Marie Sklodowska-Curie Grant Agreement No. 766181, project DeLIVER, and the European Innovation Council Pathfinder Open project DeLIVERy, grant agreement no. 101046928. P.McC. also acknowledges funding through NFR PACA-Pill, grant no. 325446.

