## Supporting Information for "Dual-color single molecule localization microscopy on transparent polymer waveguide chips"

November 2022

### 1 3D files for chip holder

We included the 3D files for the waveguide chip holder and the holder-to-stage adapter for anyone to use and modify. The .par files can be opened and modified in CAD programs such as Solid Edge (Siemens). The .stl files can be directly imported into Prusa Slicer (Prusa Research) (see Figure 1 below) for 3D print. Our parts were printed on an Original Prusa i3 MK3 printer with the settings 0.10 mm Detail MK3 for height and 0.2 infill using polylactic acid (PLA) as filament. After the chip holder is finished, it is advisable to use a scalpel or knife to remove any filament or support that might be blocking the slide of the holder and to first use a regular 24 mm coverslip to make sure that the glass slides into place on the holder.

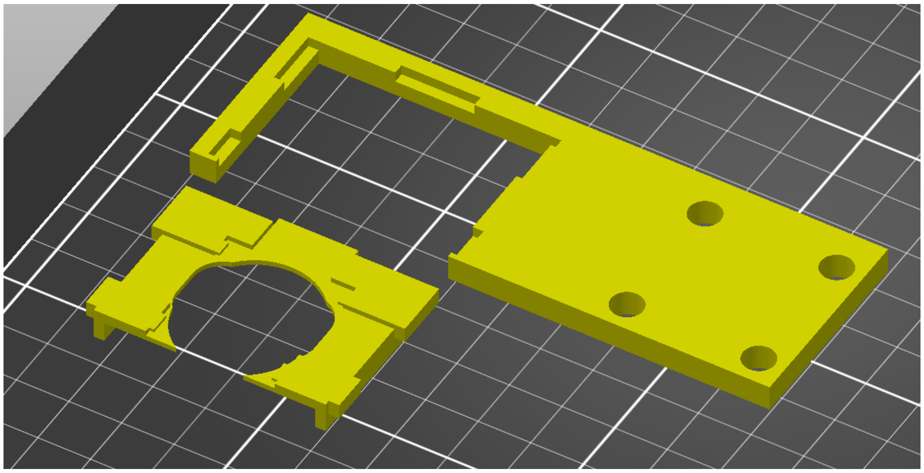

Figure 1: The sketch of the chip holder and the holder-to-stage adapter (.stl files) when imported into Prusa Slicer.
